# Combination Treatment with Intravesical Interferon-Alpha Gene Therapy and Oral Pan-ErbB Receptor Family Blocker Improves Survival in Mice with Bladder Cancer

**DOI:** 10.64898/2026.01.23.701130

**Authors:** Akshay Sood, Alberto Martini, Jan K. Rudzinski, Come Tholomier, Roberto Contieri, I-Ling Lee, Nigel R. Parker, Seppo Yla-Herttuala, David J. McConkey, Colin P.N. Dinney, Sharada Mokkapati

## Abstract

**Purpose:** Intravesical interferon-alpha (IFNα) gene therapy has shown promise in treating BCG-unresponsive non-muscle invasive bladder cancer (NMIBC). Ongoing work in our lab aims to further improve its treatment efficacy by identifying resistance mechanisms and deploying targeted combination treatment strategies.

**Experimental design:** We performed end-tumor RNA-seq analysis of MB49 murine tumors treated with IFNα gene therapy, identifying the ErbB pathway as a resistance mechanism. We consequently hypothesized that a combination treatment involving an ErbB pathway blocker and IFNα could yield improved outcomes. MB49 cells were treated *in vitro* with lentiviral IFNα (LV-IFNα) gene therapy, with/without Afatinib, a pan-ErbB inhibitor, and cell viability and migration assays were performed. Next, *in vivo* studies were conducted in the syngeneic MB49 orthotopic murine bladder cancer model. The mice were randomized into 5 treatment groups (n=10 each): saline (Ctrl), LV-Ctrl, oral Afatinib monotherapy, intravesical LV-IFNα monotherapy, and the experimental intravesical LV-IFNα + oral Afatinib combination therapy. Overall survival (OS) and drug toxicity were assessed.

**Results:** Combination therapy significantly reduced MB49 cell viability *in vitro* compared to all other treatment conditions (mean relative ATPase activity at 72 h for the combination treatment was 4%, compared to 100%, 26%, and 28% for Ctrl, LV-IFNα, and Afatinib, respectively, p<0.001). This additive effect on cell viability appeared to be driven by a combination of early-cytostatic and late-cytolytic effects. The combination treatment also markedly inhibited cell migration (mean migrated cells/10x Boyden chamber assay at 36 h were: 92.3 for the combination therapy and 631.0, 600.4, and 270.3 for Ctrl, LV-IFNα, and Afatinib, respectively, p<0.001). Finally, the *in vivo* studies demonstrated improved OS with combination therapy (median OS was 49 d in the combination group vs 15, 29, and 26 d in Ctrl, LV-IFNα, and Afatinib groups, respectively, Log-rank p<0.001). No mice in the combination therapy group died of drug toxicity.

**Conclusions:** Our preliminary findings suggest that the ErbB pathway may serve as a clinically significant resistance mechanism to intravesical IFNα gene therapy, and when targeted concurrently, may improve treatment efficacy.

## INTRODUCTION

The ErbB family of receptors plays a crucial role in several key cellular homeostatic functions such as cell growth, proliferation, differentiation, apoptosis, and motility, which, when dysregulated, can promote tumorigenesis ^1, 2^. The recently concluded TCGA project, which profiled 412 high-grade bladder tumors, also demonstrated the importance of ErbB pathway in bladder cancer pathogenesis — the EGFR, ERBB2, and ERBB3 genes were among the top 10 most frequently altered genes in patients with urothelial carcinoma of the bladder ^3^.

Nadofaragene firadenovec is an intravesically-administered, replication-deficient adenoviral vector delivered interferon-alpha (IFNα) gene therapy that has demonstrated promising results in the treatment of patients with BCG-unresponsive non-muscle invasive bladder cancer (NMIBC) ^4–6^ and was granted the FDA approval for this indication in 2022. Of the 151 patients studied in a phase-III clinical trial, 59.6% achieved complete pathologic response at the initial 3-month assessment, with 51.1% maintaining a durable response at 12 months ^5^. Although the results of this trial have been encouraging, ongoing research in our laboratory is focused on further enhancing the treatment efficacy and durability. Strategies have included biomarker-driven candidate selection ^7^, development of innovative drug delivery vectors ^8^, and identification of novel resistance mechanisms for targeted combination therapies. The current study investigates the last strategy, whereby we identified the ErbB pathway as a significant resistance mechanism to intravesical IFNα gene therapy ^8^ and sought to test the combined effect of intravesical IFNα gene therapy and an oral pan-ErbB receptor blocker on survival and tumor progression outcomes in the MB49 syngeneic orthotopic murine model of bladder cancer *in vivo* and in MB49, BBN975, and UPPL1541 murine bladder cancer cell lines *in vitro*.

## MATERIALS AND METHODS

### Cell lines

Mouse urothelial carcinoma cell line MB49/GFP-luciferase was a generous gift from Dr. Robert Svatek (The University of Texas Health, San Antonio, TX). The BBN975, derived from p53+/− mice treated with BBN (N-butyl-N-(4-hydroxybutyl)nitrosamine) and UPPL1541, derived from UPII PTEN/p53 null mice cell lines were a kind gift from Dr. William Kim (University of North Carolina, Chapel Hill, NC). Cells were grown in minimum essential medium (MB49) or Dulbecco’s modified Eagle medium (BBN975 and UPPL1541) supplemented with 10% of fetal bovine serum (FBS) and 1% penicillin-streptomycin. We interrogated these three cell lines as they encompass the molecular and physical phenotypes encountered clinically in bladder cancer ^9, 10^.

### Viral vectors

Viral vectors were provided by FKD Therapies, University of Finland (LV-Control and LV-IFNα). Transduction of mouse bladder cell lines using these vectors has been described by our laboratory previously ^8^. Briefly, for viral transduction, cells were seeded in 6-well culture dishes at varying densities: MB49 at 50,000, BBN975 at 75,000, and UPPL1541 at 75,000 cells/well. After overnight attachment, media was replaced with polybrene containing media (4 μg/mL) and viral particles were added at a desired MOI.

### Cell viability and migration assays

Cells were plated in 96-well plates at 1,000 cells/well for viability assays and in 6-well plates at 30,000 cells/well for apoptosis assays. After overnight incubation, viral vectors were added to triplicate wells at MOI of 2 and the drugs were added at their respective IC50 concentrations: MB49 0.10 μM, BBN975 0.05 μM, and UPPL1541 0.15 μM. Media containing each drug/vector was replaced every other day and cells were collected on days 1 and 3 of treatment. CellTiter-Glo luminescent method (Promega G7572) was used to determine cell viability. Annexin V allophycocyanin and propidium iodide staining (Thermo Fisher Scientific 88-8102-72) was used to determine percentage of apoptotic cells via flow cytometry. DNA content-based cell cycle analysis was also performed via flow cytometry. Cell-cycle distribution was calculated after appropriate gating of cell populations in FL-2-Area vs FL-2-Width plot of propidium iodide fluorescence. For cell migration and invasion assays, serum starved MB49, BBN975, and UPPL1541 cells (150,000 cells/well) were seeded in triplicates in the upper chamber of a Boyden Transwell chamber in 500 μL of serum-free media, separated from the chemoattractant (10% FBS supplemented media) by 8 μm pore size PCTE-PVP-free membrane pre-coated with growth-factor reduced (GFR) Matrigel (2.5 mg/mL [Cornig 354480]). After 36 hours, cells invading through the lower chamber were fixed in 4% paraformaldehyde and quantified using a 4-quadrant approach (10x magnification) via the NIH ImageJ software ^11^. All experiments had at least three biological replicates.

### In vivo syngeneic orthotopic murine model

All animal experiments were conducted in compliance with the Institutional Animal Care and Use Committee (IACUC) at the MD Anderson Cancer Center, TX. On day 0, female C57Bl/6 mice, aged 6-8 weeks, were anesthetized with isoflurane. Urethral catheterization was performed using a 20G angiocatheter. Bladders were emptied, flushed with PBS, and drained. 100 μL of PLL (0.01 μg/mL) was instilled into the bladder for 15 minutes. After draining the bladder, 25,000 MB49/GFP-luciferase cells diluted in 100 μL of hank’s balanced salt solution (HBSS) were instilled and left to dwell for 30 minutes. Following 4 days of recovery, mice were imaged using the IVIS Spectrum *In vivo* Imaging System and Living Image Software (Perkin Elmer). Mice were randomized into 5 treatment groups (n=10 animals/group) stratified by tumor uptake to assure equal baseline tumor burden distribution. Treatment and control arms consisted of saline control (Ctrl), LV-Ctrl (3 x 10^7^ virus particles in 100 μL PBS x one-time 40-minute instillation), Afatinib monotherapy (5 mg/kg daily oral gavage with rest on weekends), LV-IFNα monotherapy (3 x 10^7^ virus particles in 100 μL PBS x one-time 40-minute instillation), and the experimental LV-IFNα + Afatinib combination therapy. Tumor growth was monitored using luciferase imaging once weekly. Animals were monitored daily, and those with a reduction in body weight of greater than 20%, significant lethargy, or hematuria were deemed moribund and euthanized as per IACUC protocol. All animals were sacrificed at 9 weeks from treatment start. Laparotomy was performed and tissue specimens were collected immediately following euthanasia. Bladders were removed and weighed. Abdominal contents were assessed for metastatic disease. Tissues for RNA sequencing were collected in liquid nitrogen and tissues for histology and immunostaining were fixed in buffered formalin. The *in vivo* experiments were conducted thrice.

### Gene expression profiling and qPCR analysis

Whole transcriptome RNAseq was performed on the Ion Gene Studio S5 (Thermo Fisher Scientific). RNA was isolated from cell lines or mouse urothelium using mirVana miRNA isolation kit (Thermo Fisher Scientific) and the manufacturer instructions. Quality of RNA was assessed using Nanodrop ND-1000 spectrophotometer and 4200 Tapestation (Agilent Technologies). Twenty nanograms of RNA was transcribed into cDNA using Ion AmpliSeq transcriptome mouse gene expression chef-ready kit (A36412). cDNA was amplified and subsequently ligated with adapters and barcodes. Purified libraries were quantified using Ion library quantitation kit (Thermo Fisher Scientific) and pooled in a set of eight followed by enrichment on IonChef (Thermo Fisher Scientific). Enriched samples were then loaded onto Ion 540 chips and run on the Ion Gene Studio S5. Primary analysis of RNA-seq data was performed using the dataset GSE205493 as reference for identifying changes in the ErbB pathway. Differential expression analysis was performed using t-tests, and the false discovery rate (FDR) was estimated using the beta-uniform mixture method ^12^.

### Reverse phase protein array analysis

Cellular proteins were denatured by 1% sodium dodecyl sulphate (SDS) (with β-mercaptoethanol) and diluted in five 2-fold serial dilutions with lysis buffer. Serial diluted lysates were arrayed on nitrocellulose-coated slides by Aushon 2470 Arrayer (Aushon BioSystems). A total of 5,808 array spots were arranged on each slide including the spots corresponding to serial diluted: 1) standard lysates and 2) positive and negative controls prepared from mixed cell lysates or dilution buffer, respectively. Each slide was probed with a validated primary antibody plus a biotin-conjugated secondary antibody. Only antibodies with a Pearson correlation coefficient between Reverse Phase Protein Array (RPPA) and western blotting of greater than 0.7 were used for RPPA. Antibodies with a single or dominant band on western blotting were further assessed by direct comparison to RPPA using cell lines with differential protein expression or modulated with ligands/inhibitors or siRNA for phospho- or structural proteins, respectively. The signal obtained was amplified using a Dako Cytomation–Catalyzed system (Dako) and visualized by 3, 3’-diaminobenzidine (DAB) colorimetric reaction. The slides were scanned, analyzed, and quantified using a customized software to generate spot intensity. Each dilution curve was fitted with a logistic model (“Supercurve Fitting” developed by the Department of Bioinformatics and Computational Biology at MD Anderson Cancer Center, TX [http://bioinformatics.mdanderson.org/OOMPA]). This fits a single curve using all the samples (i.e., dilution series) on a slide with the signal intensity as the response variable and the dilution steps as the independent variables. The fitted curve is plotted with the signal intensities – both observed and fitted – on the y-axis and the log2-concentration of proteins on the x-axis for diagnostic purposes. The protein concentrations of each set of slides were then normalized for protein loading. Correction factor was calculated by 1) median-centering across samples of all antibody experiments and 2) median-centering across antibodies for each sample.

### Statistical methods

GraphPad Prism 10 Software was used for statistical analyses. Multiple group comparisons were made using two-way ANOVA or Kruskal-Wallis tests for continuous data. Chi-square test was used for categorial data. The Kaplan-Meier method using Log-rank test was utilized to perform the survival analyses. A p <0.05 was considered statistically significant.

## RESULTS

### ErbB pathway activation acts a resistance mechanism to LV-IFNα therapy

We performed post-hoc end-tumor RNA-seq analysis of MB49 murine tumors treated with LV-IFNα and LV-Ctrl (as part of a prior study) ^8^, and demonstrated a 4-to-6-fold upregulation of the ErbB pathway in treatment-resistant end-tumors. Specifically, the absolute mRNA expression levels for EGFR, ERBB2, and ERBB3 in these treatment arms were 919.6 vs 171.4, 662.7 vs 119.0, and 3,033.9 vs 679.4, respectively, p<0.001 (refer to **Figure 1a**). Based on these observations and prior investigations demonstrating high prevalence of ErbB mutations in bladder cancer ^3, 13^, we aimed to test the *in vitro* and *in vivo* efficacy of LV-IFNα + Afatinib combination therapy.

**Figure 1:**
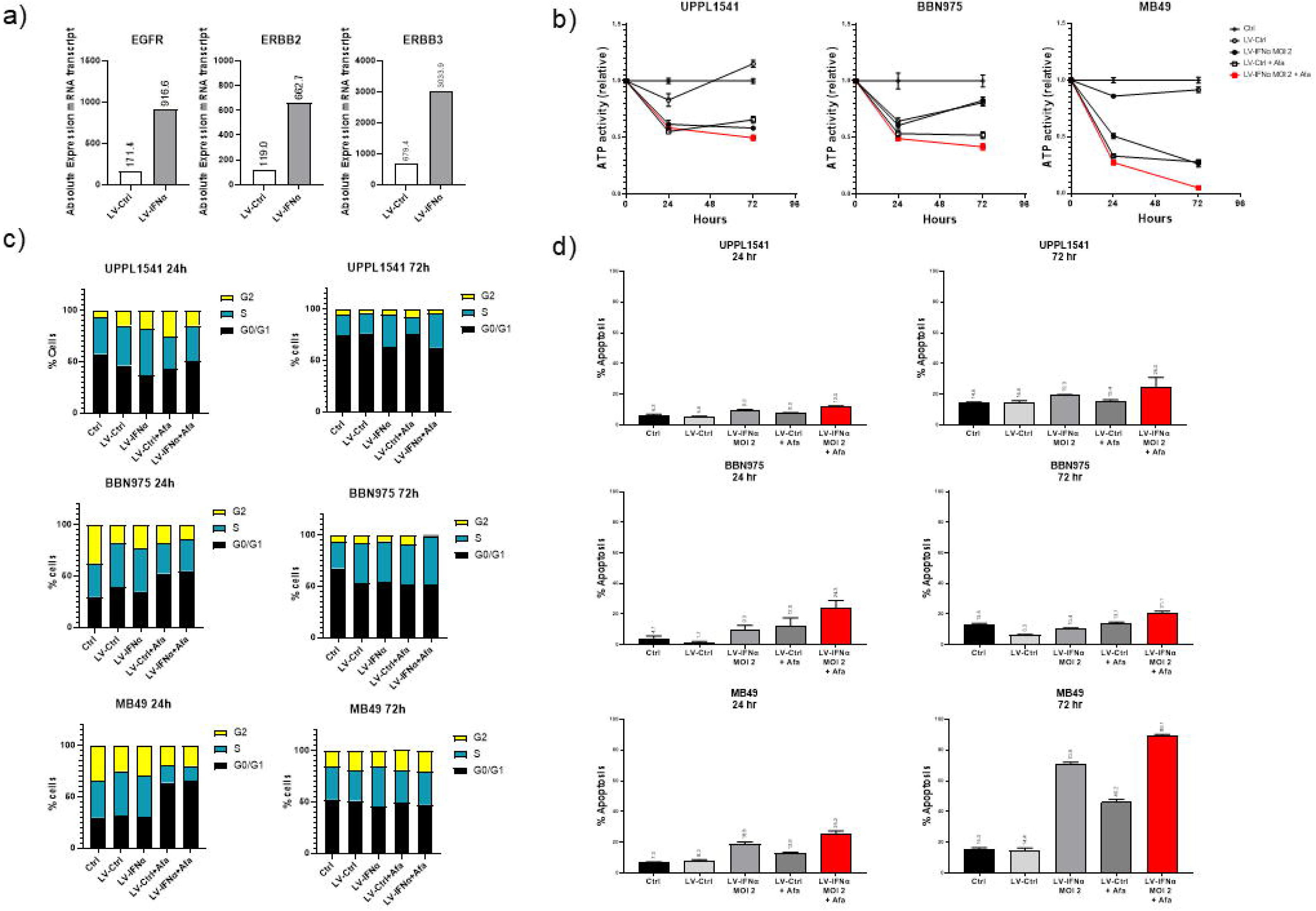
**(a)** RNA-seq analysis demonstrating EGFR, ERBB2, and ERBB3 upregulation in *in vivo* MB49 end-tumors following LV-IFNα treatment; **(b)** *In vitro* cell viability assay; **(c)** *In vitro* cell cycle (G0/G1 arrest, S phase, G2 phase) analysis at 24 h and 72 h; **(d-e)** *In vitro* apoptosis assay at 24 h and 72 h

### LV-IFNα + Afatinib combination therapy reduces cell viability and exerts additive effect on apoptosis in the MB49 murine cell line

*In vitro* testing showed that combination therapy significantly decreased the viability of MB49 cells compared to all other treatment conditions. The mean relative ATPase activity at 72 h was 4% for the combination treatment, contrasting with 100%, 96%, 26%, and 28% for the Ctrl, LV-Ctrl, LV-IFNα, and Afatinib arms, respectively, p<0.001 (**Figure 1b**). These findings were not observed in the UPPL1541 and BBN975 cell lines where combination treatment did not significantly affect cell viability compared to monotherapy (see **Figure 1b**). This additive effect on cell viability in MB49 cells appeared to be driven by a combination of early-cytostatic and late-cytolytic effects. At the 24 h mark, 65.3% of cells in the combination treatment arm were found to be in G0/G1 arrest, compared to 29.7% in the Ctrl group. This cytostatic effect was likely facilitated by the rapid onset of Afatinib’s anti-ErbB activity, as evidenced by cell-cycle arrest in the Afatinib monotherapy arm as well (**Figure 1c, bottom panel**). At the 72 h time point, cytostasis was similar across all treatment arms, however, the rate of apoptosis was significantly increased in the combination therapy arm compared to other treatment arms (89.1% in combination arm vs 70.9% and 46.2% in LV-IFNα and Afatinib arms, respectively, p<0.001) and the 24 h assessment (89.1% vs 25.2%, **Figures 1d-e, bottom panel**).

### LV-IFNα + Afatinib combination therapy reduces cell migration in all 3 murine cell lines

The combination treatment markedly inhibited cell migration and invasion across all three cell lines. The mean number of migrated MB49 cells/10x Boyden chamber assay was 92.3 for combination therapy and 631.0, 675.5, 600.4, and 270.3 for Ctrl, LV-Ctrl, LV-IFNα, and Afatinib, respectively, p<0.001 as measured at the 36 h mark (**Figure 2**). Similar results were observed for the UPPL1541 and BBN975 cell lines.

**Figure 2:**
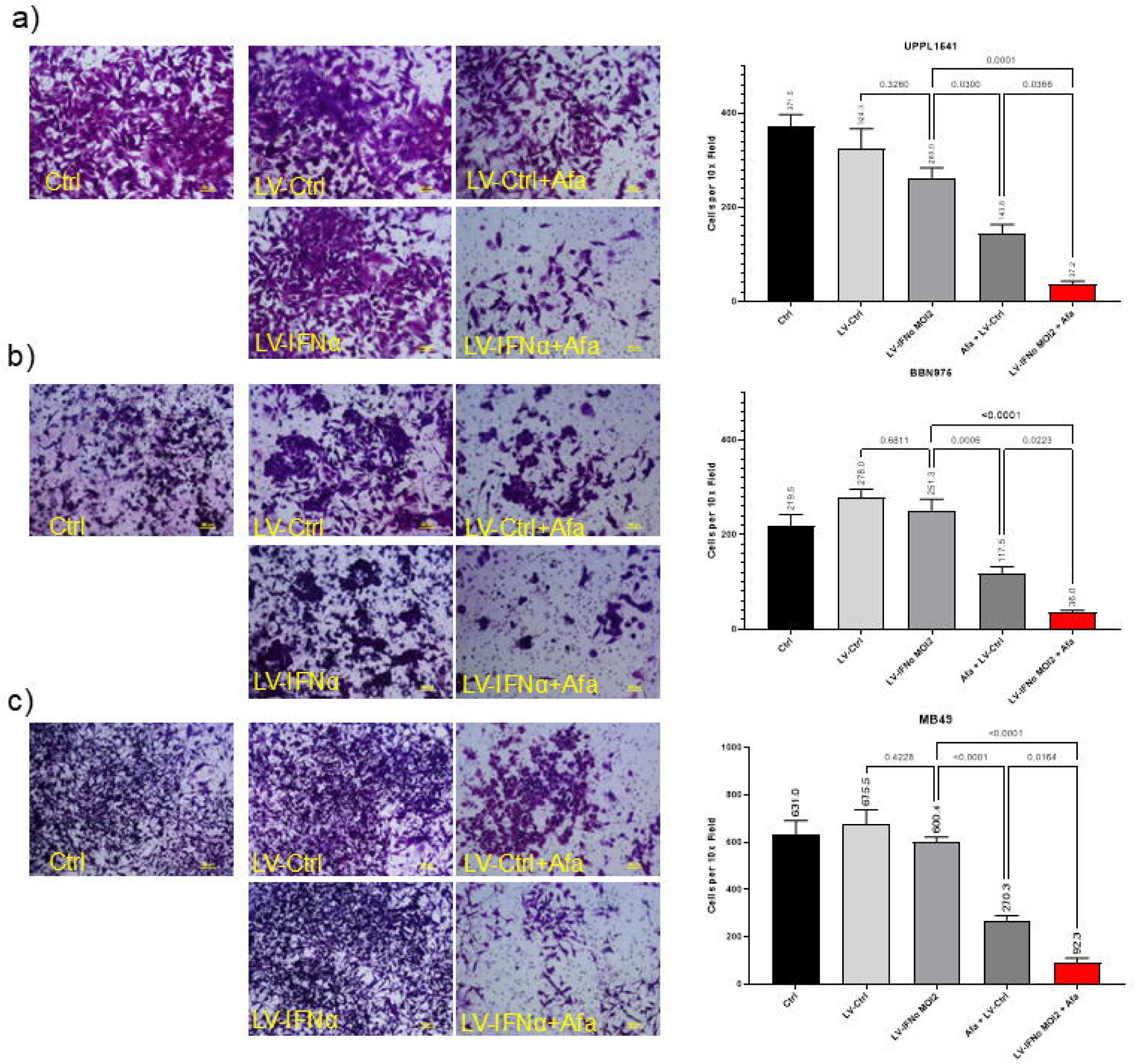
*In vitro* Boyden chamber cell migration and invasion assay at 36 h in **(a)** UPPL1541, **(b)** BBN975 and **(c)** MB49 cells treated with Ctrl, LV-Ctrl, LV-IFNα, LV-Ctrl+Afa and LV-IFNα+Afa.

### The efficacy of combination therapy correlates with the cell line’s dependence on ErbB pathway as an escape mechanism following LV-IFNα treatment

The baseline expression levels of EGFR, ERBB2, and ERBB3 mRNA transcripts in the MB49, BBN975, and UPPL1541 cell lines correlated with the observed efficacy of Afatinib monotherapy. For example, the UPPL1541 cell line, which displayed low baseline levels of ErbB receptors (**Figure 3a**), showed minimal responsiveness to Afatinib. In contrast, both BBN975 and MB49 cell lines responded well. This is in line with previous work on anti-ErbB therapy and ErbB expression in solid human tumors ^14^. The success of combination therapy, however, appeared to be tied to the reliance of each cell line on the ErbB pathway as a compensatory mechanism following IFNα therapy. As illustrated in **Figure 3b**, MB49 cells exhibited an increase in the expression of EGFR (1.8-fold), ERBB2 (1.5-fold), and ERBB3 (3.5-fold) mRNA transcripts post-LV-IFNα treatment, while the other cell lines did not. These observations were confirmed on RPPA proteomic analysis (see **Figure 3c**). Additional proteomic analysis of the MB49 cell line further showed that LV-IFNα exposure also triggered an activation of the ErbB pathway, as evidenced by enhanced phosphorylation of the EGFR-pY1173, HER2-pY1248, and HER3-pY1289 receptors and the downstream effectors (pS6-240_244 and pS6-235_236, as shown in **Figure 3d, left panel**). Afatinib therapy effectively inhibited this activation (also depicted in **Figure 3d, left panel**). Finally, Afatinib synergistically enhanced the response to LV-IFNα therapy, as indicated by increased levels of TRAIL, caspase-3, and caspase-8 — the key mediators of IFNα response (refer to **Figure 3d, middle panel**). These mechanistic insights align with the distinct tumor cell growth responses observed across the three cell lines (shown in **Figure 1b**), supporting Afatinib’s selective mechanism of action in cells that utilize the ErbB pathway as an escape mechanism.

**Figure 3:**
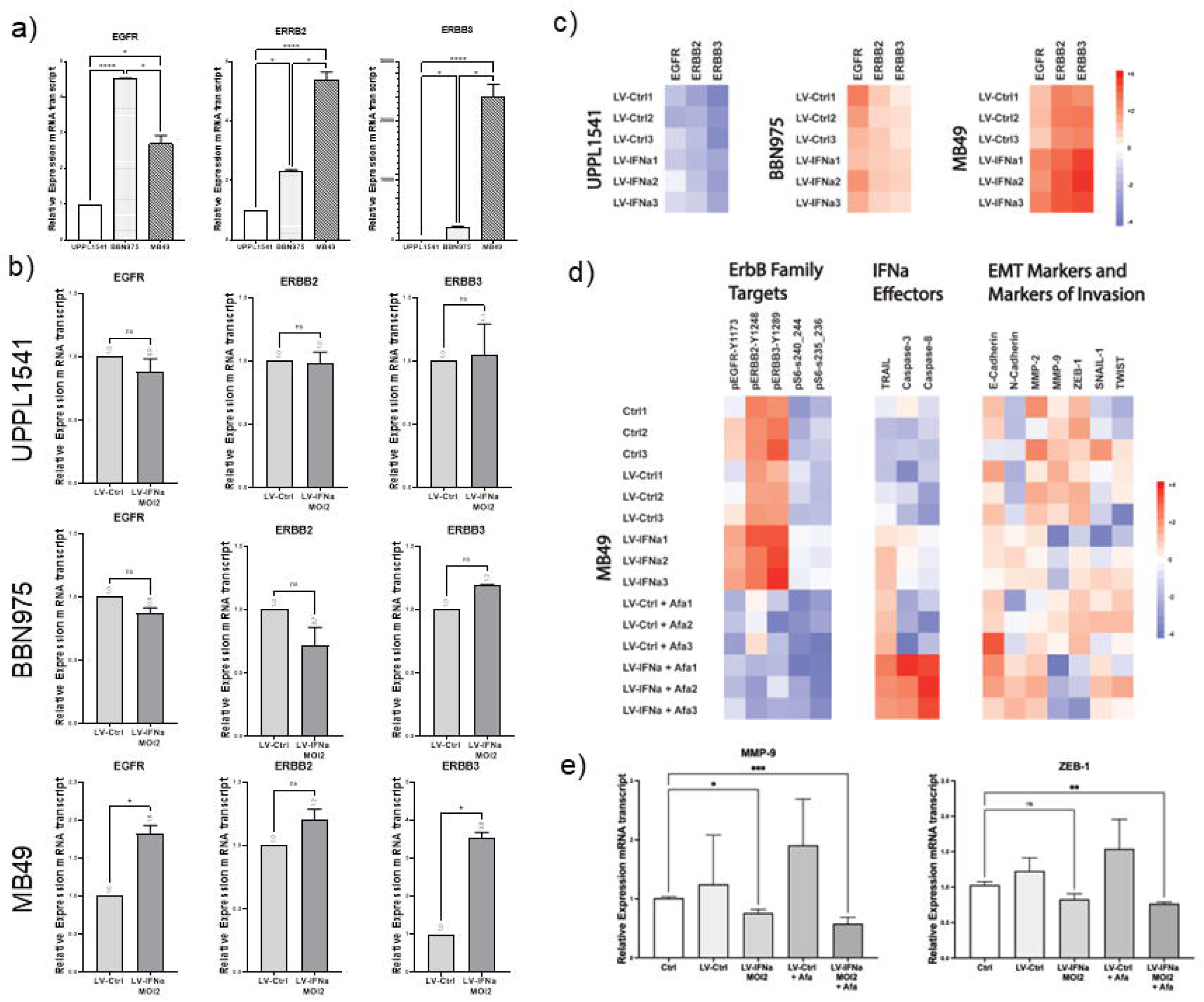
**(a)** Baseline mRNA expression of EGFR, ERBB2, and ERBB3 across the 3 cell lines; **(b)** Effect of LV-IFNα treatment on mRNA expression levels of EGFR, ERBB2, and ERBB3 across the 3 cell lines; **(c)** RPPA analysis of the effect of LV-IFNα treatment on protein-level expression of EGFR, ERBB2, and ERBB3 across the 3 cell lines; **(d)** RPPA analysis on the effect of combination treatment on ErbB, IFNα, and EMT pathways in the MB49 cell line; **(e)** RNA-seq analysis demonstrating suppression of MMP-9 and ZEB-1 with combination treatment in the MB49 cell line

### Suppression of cell invasion is likely mediated by the inhibition of MMP-9, ZEB-1, and membranous EGFR, and does not parallel the cytostatic effect of Afatinib

Preliminary findings from protein and RNA-seq analyses indicated that the suppression of MMP-9 ^15^ and ZEB-1 ^16^ likely played a role in the diminished invasive potential of MB49 cells treated with the combination therapy (refer to **Figures 3d, right panel and 3e**). Other factors such as E-cadherin, N-cadherin, MMP-2, SNAIL-1, and TWIST were also tested and did not appear to be altered by the combination therapy. MMP-9 (but not ZEB-1) was also suppressed in UPPL1541 and BBN975 cell lines [data not shown]. However, MMP-9 and ZEB-1 must work alongside other mechanisms that reduce the invasive potential, as the cells treated with Afatinib monotherapy also showed significant reduction in invasion compared to the control groups (**Figure 2**), but these cells did not demonstrate any differences in expression levels of MMP-9 and ZEB-1 (**Figure 3d, right panel**). Previous research from our laboratory has shown that a significant pool of activated EGFR is localized to the dynamic F-actin containing structures located at focal cellular motility points (such as ruffles) and is inhibited by ErbB inhibitors in a cell type agnostic fashion ^17^. Taken together, these findings suggest that the additive negative effect of combination treatment seen on motility in the Afatinib + LV-IFNα arm is likely mediated by MMP-9/ZEB-1 inhibition and ErbB receptor dephosphorylation.

### LV-IFNα + Afatinib combination therapy reduces tumor burden and improves survival

The *in vivo* studies (see **Figure 4a**) demonstrated improved OS with combination therapy. The median OS was 49 d in the combination group vs 15, 14, 29, and 26 d in the Ctrl, LV-Ctrl, LV-IFNα, and Afatinib groups, respectively, Log-rank p<0.001 (**Figure 4b**). The combination therapy group also had significantly higher mouse weights (surrogate for overall health status) when compared to other groups (**Figure 4c**) while the bladder weights (surrogate for local tumor burden) was significantly lower in the combination group (**Figures 4d**). No mice in the combination therapy group died of drug toxicity.

**Figure 4:**
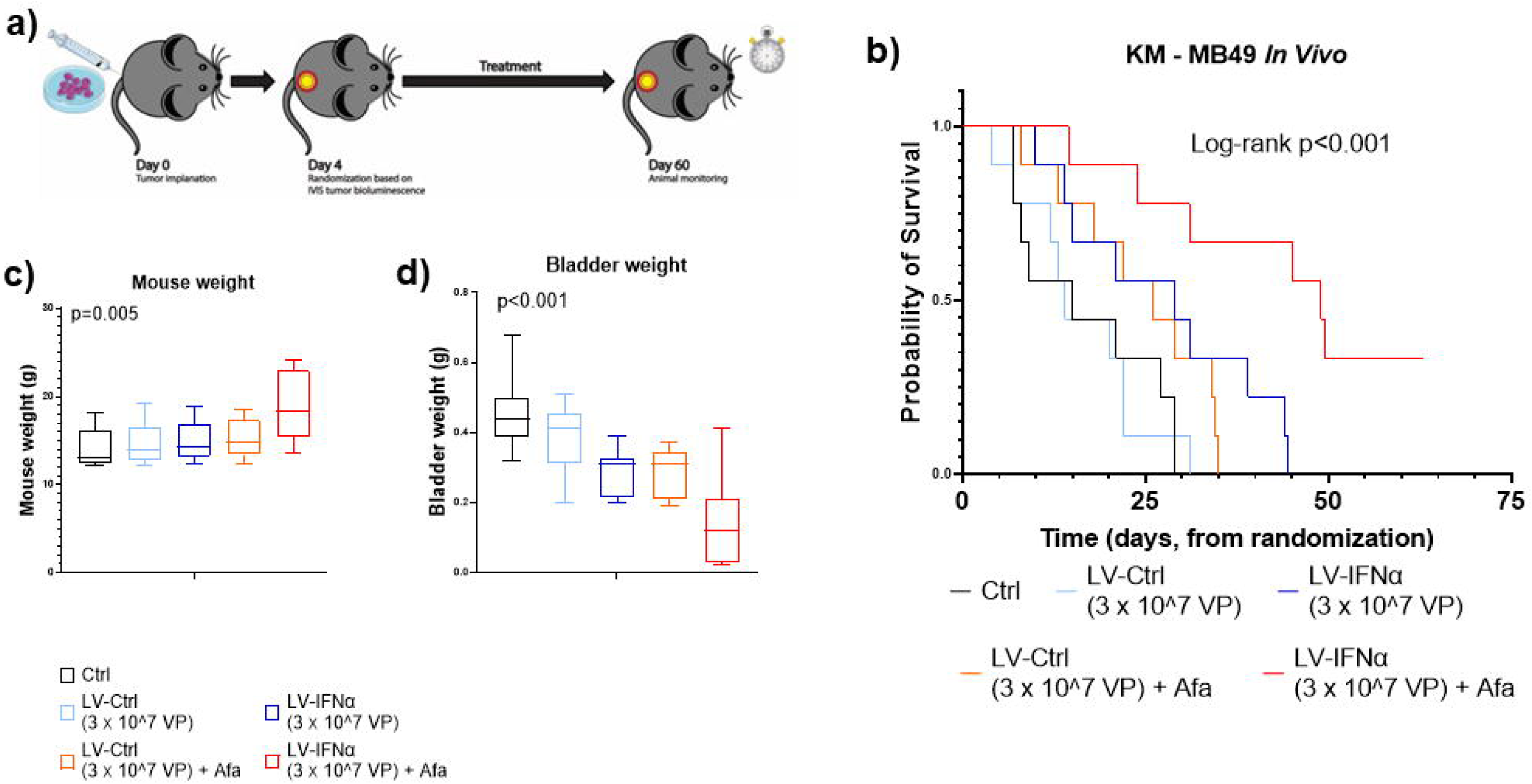
**(a)** Schematic of *in vivo* experiments in the MB49 syngeneic orthotopic bladder cancer model; **(b)** Kaplan-Meier plot of overall survival in the 5 treatment arms; **(c)** Total body weight at study end or euthanasia [surrogate for overall health status]; **(d)** Bladder weight at study end or euthanasia [surrogate for local tumor growth].

## DISCUSSION

BCG-unresponsive NMIBC is an inherently resistant cancer. Radical cystectomy remains the standard-of-care but is associated with significant morbidity and long-term quality of life considerations. Understandably, many patients are reluctant to proceed with radical cystectomy, so the identification of well-tolerated alternatives to radical cystectomy that provide durable clinical responses has become a major research focus for NMIBC.

Over the past several decades, numerous novel drugs have been evaluated for this indication including nadofaragene firadenovec ^5^, cretostimogene grenadenorepvec (CG0070) ^18^, pembrolizumab ^19, 20^, gemcitabine/docetaxel ^21, 22^, and N-803/BCG ^23^, among multiple others in trials currently ^24^. These drugs have demonstrated variable successes, but those rigorously tested in prospective trials typically exhibit an 18-month complete response rate ranging between 20-30%. The high rate of early post-treatment resistance highlights the urgent need for strategies to improve the durability and efficacy of current therapies. The results of our study underscore the potential of combining intravesical IFNα gene therapy with Afatinib, an FDA-approved oral pan-ErbB receptor blocker for the treatment of patients with BCG-unresponsive NMIBC in whom ERB pathways are activated by intravesical IFNα gene therapy. Our results show that this combination therapy significantly inhibits tumor growth (by >95%), retards cellular invasion (by >80%), and improves survival (by >300%) in a syngeneic orthotopic mouse model, highlighting its potential as a viable treatment strategy. Specifically, the efficacy of Afatinib in this combination underscores the critical role of the ErbB signaling pathway in bladder cancer pathogenesis, consistent with findings from the TCGA project which identified EGFR, HER2, and HER3 among the most frequently altered genes in high-grade bladder tumors. Our study also aligns with recent pharmacological advances in metastatic bladder cancer which have demonstrated the potential of targeting ErbB pathway in bladder malignancy via biomarker profiling of patients and specifically using either dual anti-HER-2 therapies or novel antibody-drug conjugate (ADC) agents such as disitamab-vedotin or trastuzumab-duocarmazine ^25^. While these drugs are currently being tested in refractory metastatic urothelial cancers, the same principles can be transferred to treatment-resistant NMIBC. Additionally, our work complements the recently opened ABLE-22 trial (NCT06545955), which aims to explore the concurrent use of nadofaragene firadenovec with pembrolizumab or gemcitabine/docetaxel based on similar pre-clinical work in murine models. These studies share the same goal of improving the efficacy of Ad-IFN gene therapy for patients with BCG-unresponsive NMIBC.

Our mechanistic findings align with what is known for anti-ErbB and IFNα therapies ^1, 2, 26, 27^, showcasing activation (or suppression) of well-known downstream pathways, explaining our phenotypic observations. Preliminary observations from our work also highlight the involvement of the EMT pathways in bladder cancer progression, for example the MMPs and ZEB-1 ^16^. While MMP-9 was only downregulated in the MB49 cell line, ZEB-1 was downregulated in all three murine cell lines in the combination treatment. ZEB-1, which targets E-cadherin repression (like many other EMT molecules), is a transcriptional regulator that has been implicated in progression of uterine and colorectal cancers ^28, 29^. While the findings from the current study must be regarded as preliminary, they establish the rationale to formally test the combined use of IFNα gene therapy and Afatinib in BCG-unresponsive NMIBC in a future clinical trial. This work builds upon our approach to enhanced intravesical IFNα gene therapy by identifying biomarkers that predict sensitivity and resistance and through the development of novel combination strategies that target resistance mechanisms.

## Acknowledgments

We would like to acknowledge and thank the various MDACC institutional core facilities for supporting this work, including the RPPA Core for Functional Proteomics (NCI grant #CA16672; NIH R50 #R50CA221675); The Advanced Flowcytometry and Sorting Facility at South Campus (NCI grant #P30CA16672); and Small Animal Imaging Core (Cancer Center Support grant #CA16672).

